# LRRK2 mutation alters behavioral, synaptic and non-synaptic adaptations to acute social stress

**DOI:** 10.1101/2020.02.25.965194

**Authors:** Christopher A. Guevara, Bridget A. Matikainen-Ankney, Nebojsa Kezunovic, Katherine LeClair, Alexander P. Conway, Caroline Menard, Meghan E. Flanigan, Madeline Pfau, Scott J. Russo, Deanna L. Benson, George W. Huntley

## Abstract

Parkinson’s disease (PD) risk is increased by stress and certain gene mutations, including the most prevalent PD-linked mutation *LRRK2*-G2019S. Both PD and stress increase risk for psychiatric symptoms, yet it is unclear how PD-risk genes alter neural circuitry in response to stress that may promote psychopathology. Here we show significant differences between adult G2019S knockin and wildtype (wt) mice in stress-induced behaviors, with an unexpected uncoupling of depression-like and hedonic-like responses in G2019S mice. Moreover, mutant spiny projection neurons in nucleus accumbens (NAc) lack an adaptive, stress-induced change in excitability displayed by wt neurons, and instead show stress-induced changes in synaptic properties that wt neurons lack. Some synaptic alterations in NAc are already evident early in postnatal life. Thus, G2019S alters the magnitude and direction of behavioral responses to stress that may reflect unique modifications of adaptive plasticity in cells and circuits implicated in psychopathology in humans.

## Introduction

Genetic and environmental factors collaborate to produce Parkinson’s disease (PD) in ways that are not fully understood. The most common genetic cause of late-onset PD is the G2019S mutation in leucine-rich repeat kinase 2 (LRRK2), which increases LRRK2 kinase activity by ∼2 fold (Jaleel et al., 2007; West et al., 2005). Both genetic and idiopathic forms of late-onset PD are diagnosed clinically by onset of motor system abnormalities that reflect degeneration of dopamine neurons in substantia nigra. There are prevalent non-motor symptoms associated with PD as well, including cognitive impairment and psychiatric symptoms such as depression (Gaig et al., 2014). These and other non-motor symptoms can first appear years earlier than motor symptoms and are debilitating, but not well understood mechanistically.

PD risk is increased by environmental stress and both PD and stress are associated with increased risk for depression. Brain circuits relevant to encoding lasting responses to stress are enriched for LRRK2 expression and in humans carrying G2019S, may develop, function and adapt to stressful experiences differently than those expressing wildtype (wt) LRRK2, but little is known about how environmental stress influences relevant brain circuits in ways that could promote early, PD-associated psychiatric symptoms. Social defeat stress in mice is a validated behavioral assay used to assess vulnerability to social avoidance and anhedonia-like behaviors, core features of human depression (Beery & Kaufer, 2015; Golden, Covington, Berton, & Russo, 2011). Here, we use mice carrying a G2019S knockin mutation in a coordinated set of behavioral, cellular and synaptic experiments to interrogate the effects of environmental stress on brain circuits relevant to human PD.

## Results

Young adult male wt and G2019S knockin mice underwent acute (1d) social defeat stress (1d-SDS) (**Fig 1A**). Subsequently, defeated mice were tested for social interaction (SI) by tracking their movement as they explored an arena in the absence and subsequent presence of a novel social target within a confined zone (the interaction zone, **Fig 1A**). When in the absence of a social target, 1d-SDS wt and G2019S mice spent comparable time exploring the interaction zone (**Fig 1B-D**) with no significant differences between genotypes in total distance traveled in the arena (**Fig 1E**). However, in the presence of a social target, 1d-SDS G2019S mice spent significantly less time in the interaction zone in comparison with 1d-SDS wt mice (**Fig 1B,C,F**), and traveled less overall in the arena (wt = 761.2 ± 80.13 cm, n= 13, G2019S= 565.1 ± 38.80 cm, n=12, p= 0.0428, F=4.621, Student’s t-test). Defeat experience was necessary for the increased social avoidance as unstressed G2019S and wt control mice spent equivalent amounts of time in the interaction zone in the presence of a social target (naïve wt mice = 31.23 ± 31.93 sec; naïve G2019 mice =27.99 ± 19.70 sec, n = 5 per group, p=0.8514, F=2.627, Student’s t-test). These data demonstrate that acute (1d) social defeat stress confers significantly greater social avoidance in G2019S mice than wt mice. The pronounced social avoidance of the G2019S mice from 1d-SDS was unexpected because G2019S mice undergoing chronic (10 day) social defeat stress (10d-SDS) are all highly socially interactive, while a significant proportion of wt mice undergoing 10d-SDS display prominent social avoidance (Matikainen-Ankney et al., 2018).

**Figure 1.**
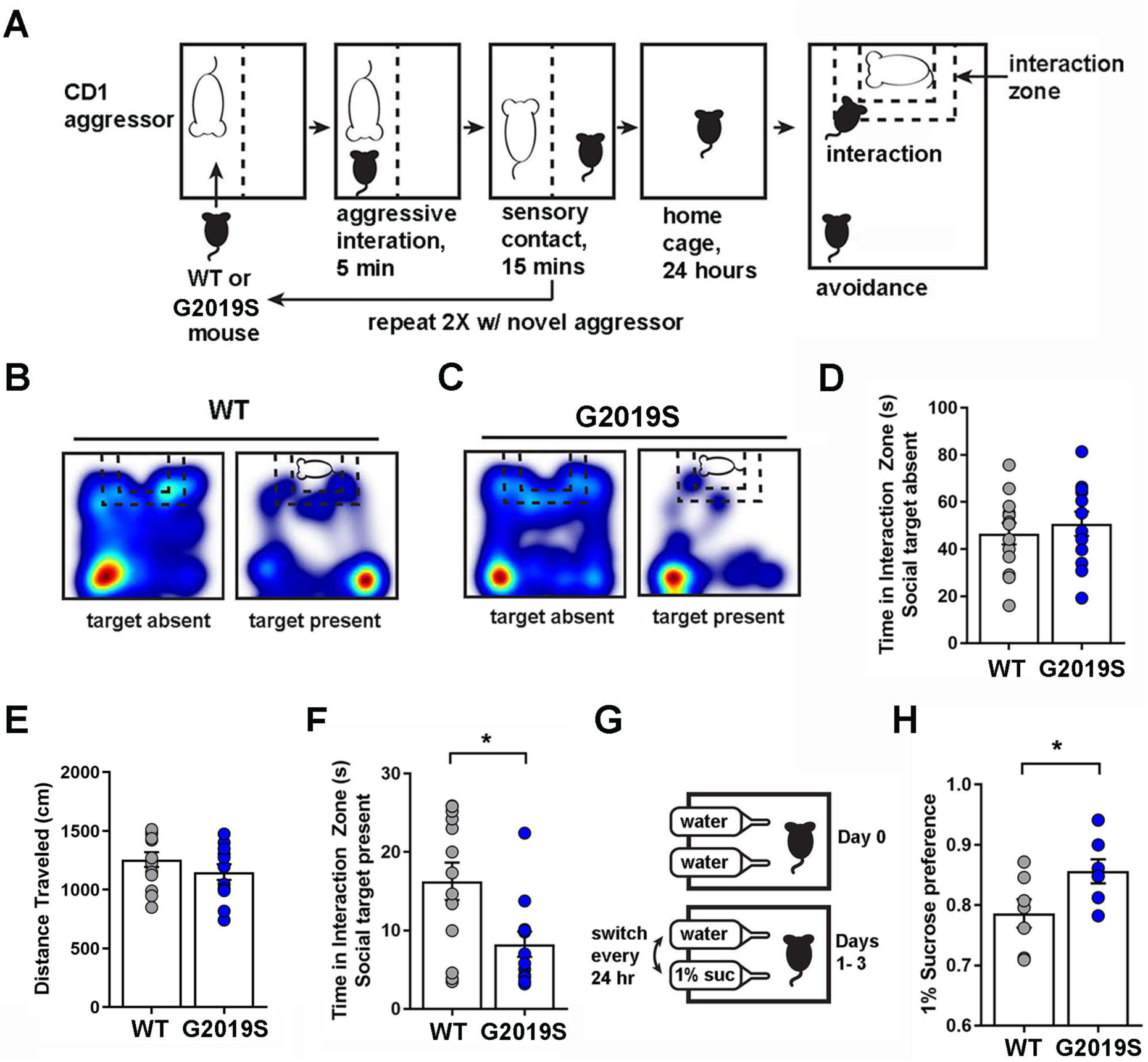
Behavioral differences between G2019S and wt mice following 1d-SDS. (**A**) 1d-SDS paradigm. (**B,C**) Representative heat maps showing movement of wt (**B**) and G2019S (**C**) mice during SI test with a novel target absent or present. Blue indicates path travelled; warmer colors indicate increased time. (**D**) Graph of time in interaction zone with no social target present (p=0.5478, F=1.186). (**E**) Total distance travelled in the arena (p=0.2593, F=1.023) (**F**) Graph of time in interaction zone with a novel social target present (p= 0.0117, F=2.404; wt (n=13 mice), G2019S (n=12 mice)). (**G**) Schematic showing 3-day sucrose preference test paradigm. (**H**) Following 1d-SDS, G2019S mice show greater sucrose consumption compared to wt (p = 0.0432, F=1.399; wt (n= 7 mice), G2019S (n =8 mice)). **D, E, F**, and **H**, Student’s t-tests.

Wildtype mice that display social avoidance following 10d-SDS also display anhedonic-like behaviors (Beery & Kaufer, 2015; Golden et al., 2011). To test the prediction that 1d-SDS-G2019S mice would therefore also display greater levels of anhedonia-like behavior, we subjected wt and G2019S mice to a 3-day sucrose-preference test post-1d-SDS (**Fig 1G**). Surprisingly, G2019S mice showed significantly *increased* average sucrose consumption compared to wt mice (**Fig 1H**), thereby revealing an unexpected uncoupling of “depression-like” and “anhedonic-like” behaviors. In the absence of defeat experience, wt and G2019S mice display similar levels of sucrose consumption (Matikainen-Ankney et al., 2018).

Following behavioral characterization, we interrogated underlying modifications to intrinsic excitability and synaptic responses in 1d-SDS-exposed mice by preparing acute slices for whole-cell patch clamp recordings from spiny projection neurons (SPNs) in the nucleus accumbens (NAc), an area rich in LRRK2 expression and known to regulate stress responses and depression-like behaviors in mice and humans (Bosch-Bouju, Larrieu, Linders, Manzoni, & Laye, 2016; Carlezon, Duman, & Nestler, 2005; Han & Nestler, 2017). We first established that in unstressed controls, there were no significant differences between genotypes in intrinsic excitability of SPNs--assessed by comparing the number of action-potentials (APs) generated in response to depolarizing current steps (**Fig. 2A,B**) and by rheobase, the amount of threshold current required to generate the first AP (**Fig. 2C**)--nor were there significant differences in interevent interval (IEI) or amplitude of spontaneous excitatory postsynaptic currents (sEPSCs) (**Fig. 2D-F**). Unexpectedly however, we found that subsequent 1d-SDS-induced cell- and synaptic adaptations differed substantially between genotypes, with wt SPNs showing significant changes in excitability and G2019S neurons showing significant changes in synaptic properties. Following 1d-SDS, wt SPNs displayed significantly increased intrinsic excitability compared to 1d-SDS-G2019S SPNs or to SPNs in unstressed controls (**Fig. 2A-C**), but no significant changes in sEPSC IEI (**Fig. 2D,E**) or amplitude (**Fig. 2D,F**). The elevation in neuronal excitability occurred without changes in resting membrane potential (p >0.99). In contrast, the intrinsic excitability of 1d-SDS-G2019S SPNs was unchanged in comparison with either wt or G2019S unstressed control SPNs (**Fig. 2A-C**). However, following 1d-SDS, G2019S SPNs exhibited a significant decrease in sEPSC IEI (**Fig. 2E**) and a significant increase in sEPSC amplitude (**Fig. 2F**), changes in synaptic properties that 1d-SDS wt SPNs lacked (**Fig. 2D-F**). These data reveal two novel findings. First, acute social stress in wt mice drives significant, presumably adaptive plasticity of intrinsic excitability of SPNs without changing their baseline synaptic properties. Two, acute stress in G2019S mice significantly alters behavioral outcomes in comparison with wt mice, fails to affect membrane excitability but produces changes in sEPSC frequency and amplitude.

**Figure 2.**
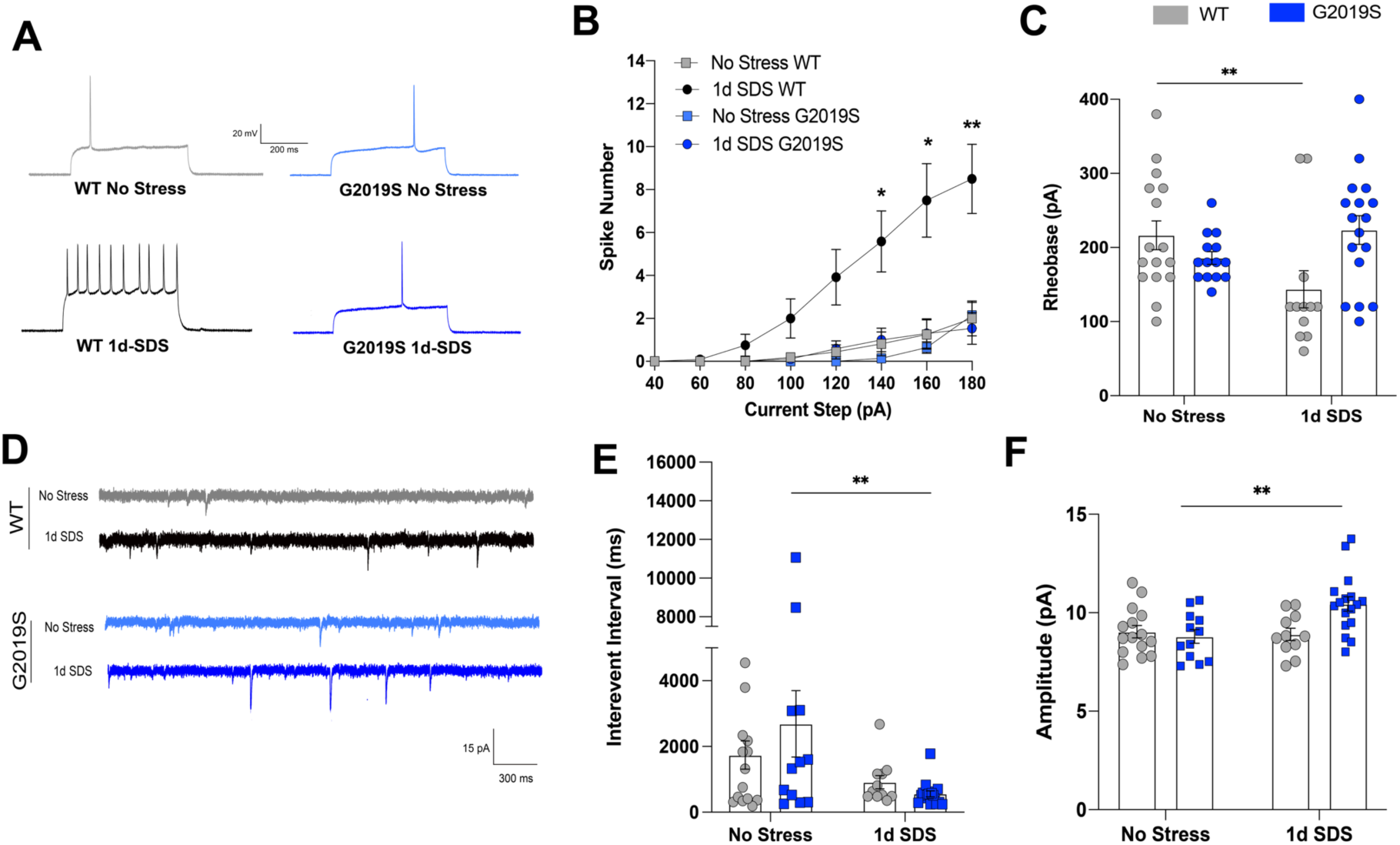
Differential adaptations in excitability and synaptic properties in G2019S and wt SPNs following 1d-SDS. (**A**) Representative traces of action potentials generated by current injection (−180 pA) into NAc SPNs, taken from mice from each behavioral condition shown. (**B**) Plot showing number of spikes elicited as a function of increasing current injected into SPNs from wt or G2019S mice from the behavioral conditions shown. Increased excitability was observed in 1d-SDS wt SPNs compared with no stress wt SPNs (at 140 pA: p=0.0189; at 160 pA: p=0.0115; at 180 pA: p=0.0064). (**C**) Graph showing average rheobase recorded from SPNs in wt or G2019S mice following no-stress or 1d-SDS conditions. wt SPNs following 1d-SDS show significantly decreased rheobase in comparison with no stress wt SPNs (p = 0.0098). (**D**) Sample traces of sEPSCs recorded from wt or G2019S SPNs for each condition. (**E**) Graph showing average interevent interval (IEI) of sEPSCs recorded from wt or G2019S SPNs. Following 1d-SDS, sEPSC IEI is significantly lower in G2019S SPNs compared to no stress G2019S SPNs (p = 0.0099). (**F**) Graph showing average amplitude of sEPSCs recorded from wt or G2019S SPNs. Following 1d-SDS, G2019S SPNs show significantly increased amplitude in comparison with G2019S no stress SPNs (p=0.0025). For all experiments, n =4-5 mice/12-17 cells per group. Repeated measures two-way ANOVA was applied to data shown in **B**. Mixed effects models with Sidak post hoc tests for multiple comparisons were applied to data shown in **C, E, F**. *p < 0.05, **p <0.01, ***p <0.001. Error bars represent S.E.M.

Because PD risk genes are carried throughout life and could be expected to influence circuit formation in NAc, we probed G2019S or wt NAc SPNs at P21 for differences in baseline synaptic properties that may already be established early postnatally. The data show that G2019S SPNs exhibited significantly greater amplitude of sEPSCs (**Fig. 3A-C**) and larger evoked AMPAR-mediated responses compared to wt (**Fig. 3D,E**), suggesting stronger glutamatergic synapses. There were no significant differences between genotypes in sEPSC IEI (**Fig. 3F**). Because stronger synapses are correlated with larger dendritic spines (Yuste & Bonhoeffer, 2001), we compared spine morphology of biocytin-filled mutant and wt SPNs following whole-cell recording. While there was no significant difference between genotypes in average spine density (**Fig 3G**), cumulative probability distribution of spine-head widths showed G2019S spines were shifted significantly towards larger values compared to wt as predicted by the larger current amplitudes (**Fig 3H,I**). Thus, structural and functional abnormalities in NAc synapses are evident during a period in which striatal circuitry can be readily and permanently modified by activity (Kozorovitskiy, Saunders, Johnson, Lowell, & Sabatini, 2012) and may impact behavioral outcomes that depend on such circuitry later in life.

**Figure 3.**
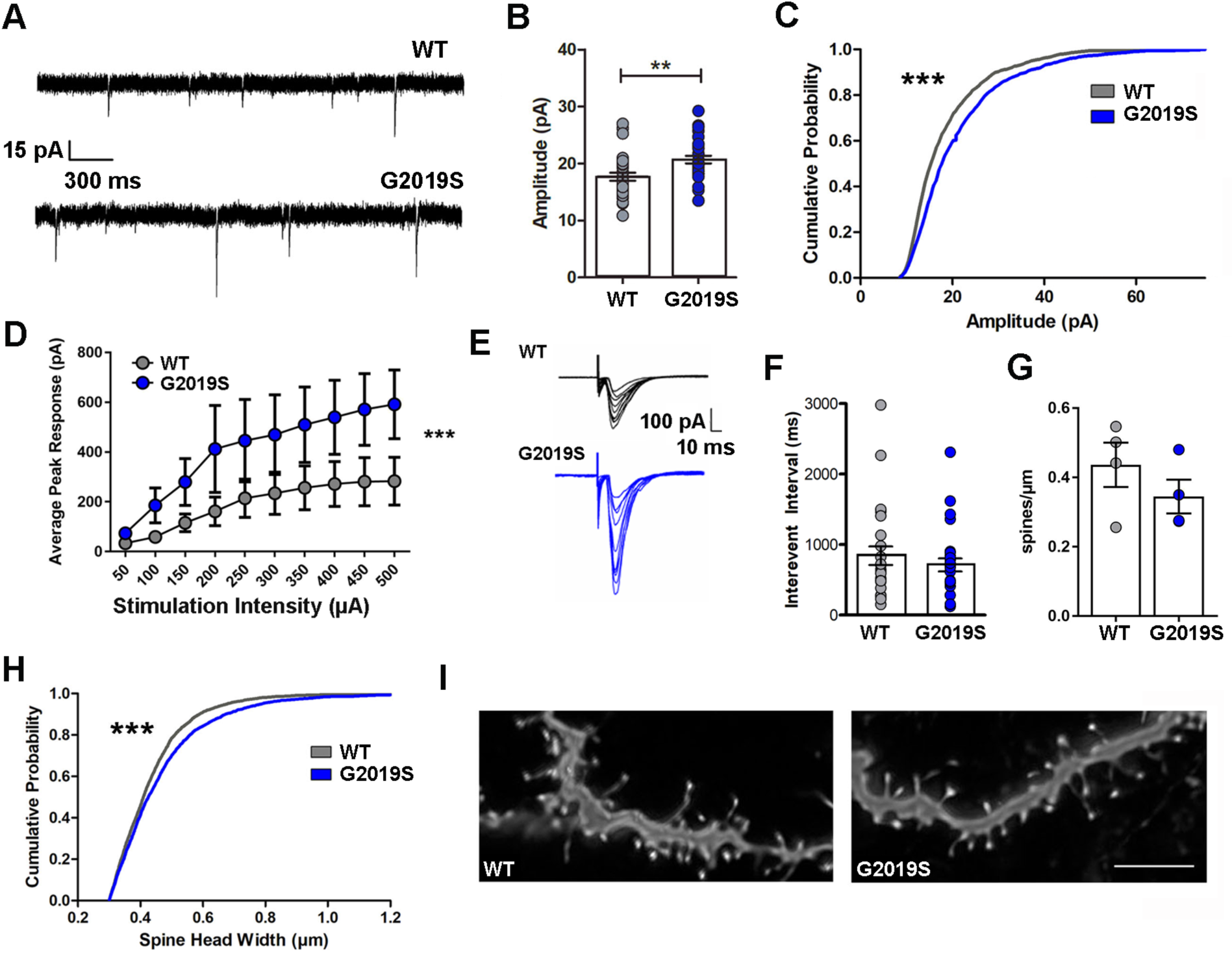
G2019S increases sEPSC amplitude and spine head-width in NAc SPNs at P21. (**A**) Sample traces of sEPSCs recorded from wt or G2019S NAc SPNs. (**B**) Graphs showing average amplitudes of sEPSCs (p= 0.0028, F=1.043. WT n=30(4), G2019S, n=30(4)). (**C**) Cumulative probability distributions of sEPSC amplitudes of the first 50 events per cell from wt or G2019S SPNs. Rightward shift is significant, p=0.0001, WT n=30(4), G2019S n=30(4). (**D**) AMPAR-current input-output curve: evoked current magnitude vs. increasing input current for wt or G2019S (n= 12(4) each group), p<0.0001 for genotype effect. (**E**) Example traces from wt or G2019S evoked AMPAR currents. (**F**) Average interevent intervals (IEI) of sEPSCs from wt or G2019S SPNs. wt, n=30(4), G2019S, n=30(4), p= 0.1047, F=2.809 and p=0.4064, F=1.893, respectively. (**G**) Graph of average spine densities per animal, wt vs. G2019S; p=0.2963, n=3-4 animals/genotype. (**H**) Cumulative probability distributions of wt or G2019S SPN spine head widths. Spine-head widths in G2019S SPNs show a significant rightward shift, p = 0.0001; n=20(4) for each group. (**I**) Examples of deconvolved (Autoquant) confocal image z-stacks (100X objective, Zeiss LSM780; Nyquist sampling) of biocytin filled, Alexa594-labeled G2019S or wt SPN dendrite segments; scale bar = 4 μm. All graphs, gray = WT; blue = G2019S; **B, F**, and **G**, Student’s t-test. **C** and **H**, Kolmogorov-Smirnov test. **D**, 2-way ANOVA.

## Discussion

Together, these data highlight three main points. First, G2019S mice respond to acute social stress in ways that are significantly different from wt mice. Moreover, the prominent social avoidance of the mutant mice after 1d-SDS was unanticipated because G2019S mice exposed to 10d-SDS are all significantly *more* socially interactive than 10d-SDS wt mice (Matikainen-Ankeny et al, 2018). Further, G2019S mice--regardless of the amount of social defeat (1d or 10d) or the degree of social interaction following defeat--display increased sucrose consumption (**Fig 1H**) (Matikainen-Ankney et al., 2018) revealing an unexpected disassociation between social interaction behavior and hedonic responses to sucrose following social stress. These differing behavioral responses require defeat experience as no differences between genotypes were evident in its absence. Thus, stress-induced behaviors in G2019S mice defy categorization as “depression-like” or “resilient-like”, and instead suggest G2019S imparts a complex set of temporally evolving behavioral responses to social stress likely reflecting both the nature and duration of the stressor. Whether this is a signature of PD vulnerability or a contributor to disease onset or progression is not known. Future studies will need to test responses to other forms and durations of behavioral stress.

Second, we found an unexpected non-synaptic adaptation to acute stress in wt SPNs that was absent in G2019S SPNs. Although no studies we are aware of have examined intrinsic excitability of SPNs following 1d-SDS, previous studies in wt mice undergoing 10d-SDS have shown that D_1_R-SPNs, but not D_2_R-SPNs, exhibit significantly increased intrinsic excitability but only in those chronic SDS mice showing prominent social avoidance (Francis et al., 2015). While such non-synaptic plasticity may be one of the earliest cellular adaptations to SDS, how such non-synaptic adaptations ultimately influence social interaction or hedonic behaviors is not clear. The complete lack of such stress-induced excitability changes in G2019S mice, coupled with modest but significant synaptic changes in sEPSC amplitude and frequency, which were lacking in wt mice, may have together maladaptively contributed to the prominent social avoidance and/or increased sucrose consumption displayed by the mutants, but this, along with potential differences between SPN subtypes, remains to be tested. While the mechanisms preventing excitability changes or promoting changes in sEPSCs are not yet known, it is plausible that altered function of the Rab family of GTPases, which are principal LRRK2 phosphotargets and important for trafficking of membrane channels and receptors (Seol, Nam, & Son, 2019; Steger et al., 2016), could underlie both synaptic and non-synaptic abnormalities observed in the mutants. The Rho family of GTPases has been implicated in excitability changes following chronic SDS (Francis, Gaynor, Chandra, Fox, & Lobo, 2019).

Third, we show that glutamatergic synaptic response strength and spine morphology of NAc SPNs were significantly different than wt already by early postnatal ages. This is particularly notable because in striatum and elsewhere, developing synaptic circuits exhibit sensitive periods during the first few postnatal weeks where altered activity persistently changes cell properties and network function (Lieberman et al., 2018; Peixoto, Wang, Croney, Kozorovitskiy, & Sabatini, 2016). This suggests G2019S co-opts synaptic circuits early in life with enduring consequences for altered stress-related responses by young adulthood. Over time this may alter synapse plasticity (Christoffel et al., 2011; Derks, Krugers, Hoogenraad, Joels, & Sarabdjitsingh, 2016; Stelly, Pomrenze, Cook, & Morikawa, 2016) and inflammatory pathways (Zhu, Klomparens, Guo, & Geng, 2019), ultimately increasing PD vulnerability.

## Methods

### Mice

*Lrrk2*-G2019S knockin mice were generated by Eli Lilly and characterized previously (Matikainen-Ankney et al., 2018; Matikainen-Ankney et al., 2016). Mice were congenic on C57Bl/6NTac background, bred as homozygotes, and backcrossed to wt C57Bl/6NTac every fourth generation to prevent genetic drift. Age- and strain-matched wildtype (wt) mice bred and raised under conditions identical to G2019S mice were used as controls. Male and female mice (aged P21) were used for electrophysiology and spine morphology using methods described in detail previously (Matikanen-Ankeny et al, 2016). Male mice (10-12 weeks old) were used for behavior. CD1 retired breeders (Charles River, Raleigh, NC) were ≥ 4 months and were screened for aggression. Animal procedures were approved by Mount Sinai’s Institutional Animal Care and Use Committee and conformed to National Institutes of Health guidelines.

### Electrophysiology

Whole-cell patch clamp recordings from spiny projection neurons (SPNs) in the NAc shell were conducted on acute coronal slices taken from unstressed wt or G2019S mice or those undergoing 1d-SDS, using methods described in detail previously (Matikainen-Ankney et al., 2018; Matikainen-Ankney et al., 2016). SPNs were identified visually and electrophysiologically (Matikainen-Ankney et al., 2016), sEPSCs were confirmed to be glutamatergic as described (Matikainen-Ankney et al., 2016).

### Behavior

For 1d-SDS, age-matched wt or G2019S male mice were subjected to brief periods of physical subordination by a larger aggressor mouse as depicted (**Fig. 1A**). Social interaction (SI) was assessed in the absence and subsequent presence of a novel social target as described (Golden et al., 2011; Matikainen-Ankney et al., 2018). Mouse movement was continuously tracked (Ethovision 5.0; Noldus). For sucrose preference, mice were given a choice of water or 1% sucrose solution as outlined in **Fig 1G**. Amount of sucrose consumed: (vol sucrose consumed/total vol liquid consumed)*100.

### Statistical Analyses

P < 0.05 was considered significant. Analyses were derived from GraphPad Software Prism (v8.2.1). Data are presented as mean values ± SEM. Numbers (n) listed as: n= number of cells (number of mice) or n= number of mice.

## References

Beery, A. K., & Kaufer, D. (2015). Stress, social behavior, and resilience: insights from rodents. Neurobiol Stress, 1, 116–127. doi:10.1016/j.ynstr.2014.10.004

Bosch-Bouju, C., Larrieu, T., Linders, L., Manzoni, O. J., & Laye, S. (2016). Endocannabinoid-Mediated Plasticity in Nucleus Accumbens Controls Vulnerability to Anxiety after Social Defeat Stress. Cell Rep, 16 (5), 1237–1242. doi:10.1016/j.celrep.2016.06.082

Carlezon, W. A., Jr., Duman, R. S., & Nestler, E. J. (2005). The many faces of CREB. Trends Neurosci, 28 (8), 436–445. doi:10.1016/j.tins.2005.06.005

Christoffel, D. J., Golden, S. A., Dumitriu, D., Robison, A. J., Janssen, W. G., Ahn, H. F., … Russo, S. J. (2011). IkappaB kinase regulates social defeat stress-induced synaptic and behavioral plasticity. J Neurosci, 31 (1), 314–321. doi:10.1523/jneurosci.4763-10.2011

Derks, N. A., Krugers, H. J., Hoogenraad, C. C., Joels, M., & Sarabdjitsingh, R. A. (2016). Effects of Early Life Stress on Synaptic Plasticity in the Developing Hippocampus of Male and Female Rats. PLoS One, 11 (10), e0164551. doi:10.1371/journal.pone.0164551

Francis, T. C., Chandra, R., Friend, D. M., Finkel, E., Dayrit, G., Miranda, J., … Lobo, M. K. (2015). Nucleus accumbens medium spiny neuron subtypes mediate depression-related outcomes to social defeat stress. Biol Psychiatry, 77 (3), 212–222. doi:10.1016/j.biopsych.2014.07.021

Francis, T. C., Gaynor, A., Chandra, R., Fox, M. E., & Lobo, M. K. (2019). The Selective RhoA Inhibitor Rhosin Promotes Stress Resiliency Through Enhancing D1-Medium Spiny Neuron Plasticity and Reducing Hyperexcitability. Biol Psychiatry, 85 (12), 1001–1010. doi:10.1016/j.biopsych.2019.02.007

Gaig, C., Vilas, D., Infante, J., Sierra, M., Garcia-Gorostiaga, I., Buongiorno, M., … Tolosa, E. (2014). Nonmotor symptoms in LRRK2 G2019S associated Parkinson’s disease. PLoS One, 9 (10), e108982. doi:10.1371/journal.pone.0108982

Golden, S. A., Covington, H. E., 3rd, Berton, O., & Russo, S. J. (2011). A standardized protocol for repeated social defeat stress in mice. Nat Protoc, 6 (8), 1183–1191. doi:10.1038/nprot.2011.361

Han, M. H., & Nestler, E. J. (2017). Neural Substrates of Depression and Resilience. Neurotherapeutics. doi:10.1007/s13311-017-0527-x

Jaleel, M., Nichols, R. J., Deak, M., Campbell, D. G., Gillardon, F., Knebel, A., & Alessi, D. R. (2007). LRRK2 phosphorylates moesin at threonine-558: characterization of how Parkinson’s disease mutants affect kinase activity. Biochem J, 405 (2), 307–317. doi:10.1042/bj20070209

Kozorovitskiy, Y., Saunders, A., Johnson, C. A., Lowell, B. B., & Sabatini, B. L. (2012). Recurrent network activity drives striatal synaptogenesis. Nature, 485 (7400), 646–650. doi:10.1038/nature11052

Lieberman, O. J., McGuirt, A. F., Mosharov, E. V., Pigulevskiy, I., Hobson, B. D., Choi, S., … Sulzer, D. (2018). Dopamine Triggers the Maturation of Striatal Spiny Projection Neuron Excitability during a Critical Period. Neuron, 99 (3), 540–554.e544. doi:10.1016/j.neuron.2018.06.044

Matikainen-Ankney, B. A., Kezunovic, N., Menard, C., Flanigan, M. E., Zhong, Y., Russo, S. J., … Huntley, G. W. (2018). Parkinson’s Disease-Linked LRRK2-G2019S Mutation Alters Synaptic Plasticity and Promotes Resilience to Chronic Social Stress in Young Adulthood. J Neurosci, 38 (45), 9700–9711. doi:10.1523/jneurosci.1457-18.2018

Matikainen-Ankney, B. A., Kezunovic, N., Mesias, R. E., Tian, Y., Williams, F. M., Huntley, G. W., & Benson, D. L. (2016). Altered Development of Synapse Structure and Function in Striatum Caused by Parkinson’s Disease-Linked LRRK2-G2019S Mutation. J Neurosci, 36 (27), 7128–7141. doi:10.1523/JNEUROSCI.3314-15.2016

Peixoto, R. T., Wang, W., Croney, D. M., Kozorovitskiy, Y., & Sabatini, B. L. (2016). Early hyperactivity and precocious maturation of corticostriatal circuits in Shank3B(-/-) mice. Nat Neurosci, 19 (5), 716–724. doi:10.1038/nn.4260

Seol, W., Nam, D., & Son, I. (2019). Rab GTPases as Physiological Substrates of LRRK2 Kinase. Exp Neurobiol, 28 (2), 134–145. doi:10.5607/en.2019.28.2.134

Steger, M., Tonelli, F., Ito, G., Davies, P., Trost, M., Vetter, M., … Mann, M. (2016). Phosphoproteomics reveals that Parkinson’s disease kinase LRRK2 regulates a subset of Rab GTPases. Elife, 5. doi:10.7554/eLife.12813

Stelly, C. E., Pomrenze, M. B., Cook, J. B., & Morikawa, H. (2016). Repeated social defeat stress enhances glutamatergic synaptic plasticity in the VTA and cocaine place conditioning. Elife, 5. doi:10.7554/eLife.15448

West, A. B., Moore, D. J., Biskup, S., Bugayenko, A., Smith, W. W., Ross, C. A., … Dawson, T. M. (2005). Parkinson’s disease-associated mutations in leucine-rich repeat kinase 2 augment kinase activity. Proc Natl Acad Sci U S A, 102 (46), 16842–16847. doi:10.1073/pnas.0507360102

Yuste, R., & Bonhoeffer, T. (2001). Morphological changes in dendritic spines associated with long-term synaptic plasticity. Annu Rev Neurosci, 24, 1071–1089. doi:10.1146/annurev.neuro.24.1.1071

Zhu, Y., Klomparens, E. A., Guo, S., & Geng, X. (2019). Neuroinflammation caused by mental stress: the effect of chronic restraint stress and acute repeated social defeat stress in mice. Neurol Res, 41 (8), 762–769. doi:10.1080/01616412.2019.1615670

